# Mechanical resistance of the environment affects root hair growth and nucleus dynamics

**DOI:** 10.1101/2022.12.15.520546

**Authors:** David Pereira, Thomas Alline, Emilie Lin, Atef Asnacios

## Abstract

Root hair (RH) cells are important for the growth and survival of seedlings. They favor plant-microbe interactions, nutrients, and water uptake. RH cells increase drastically the surface of exchange of the root system with the surrounding environment. To be able to invade the soil, RH cells have to penetrate a dense and porous medium exhibiting a variety of physical properties. The soil’s physical properties, such as mechanical resistance, impact the growth and survival of plants. Consequently, studying the effect of soil resistance on the growth of RH is essential to improve our understanding of plant growth. Here we investigate the effect of the mechanical resistance of the culture medium on RH-physical and phenotypical parameters such as length, time, and speed of growth. We also analyze the impact of the environment on the positioning, and movement of the nucleus inside the growing cells. To do so, Arabidopsis Thaliana seedlings were cultured in a custom-made microfluidic-like system, in solid media with agar concentrations ranging from 0.5% to 1.25%. We show that the time of growth of RH cells is independent of the mechanical resistance of the surrounding environment, while the RH speed decreases when the mechanical resistance increases. As a consequence, the RH cells are shorter in stiffer environments. Moreover, we show that the speed of the nucleus adapts to the mechanical resistance of the environment and follows the same trend as the average speed of the RH tip. Eventually, during RH growth, the nucleus-to-tip distance was found to decrease when the stiffness of the environment was increased, indicating mechanotransduction from the cell surface to the nucleus.

## Introduction

Roots are important structure for plant anchorage and water uptake(1)(2). The external layer of the root in contact with the surrounding environment is called the epidermis. This layer is composed of atrichoblast and trichoblast cells, the latter differentiates in tubular structure called root hair cells (RHs). They are tip growing cells of 10 μm in diameter and can be few millimeters long(3).

They are important for the plant soil interaction and nutrients uptake. Indeed, these tubular structures can increase drastically the surface of exchange of the root with the surrounding environment (up to 2-fold)(4). In addition, they are important for root-substrate “cohesion” and soil erosion(5), but also for root anchoring and penetration(6). These cells have to evolve in heterogeneous environment presenting a variety of chemical and mechanical constraints. Depending on the soil strength (moistening, density, grain size and shape) the RH cells have to face a slightly or highly growth-resisting environment.

A key component of the root hair cell’s interaction with the surrounding environment is the cell wall (CW). It is a rigid structural component that acts like a barrier to maintain cell integrity by preventing cell from over-expansion/bursting under turgor pressure and protect it from damages due to mechanical stresses. RH cell growth is achieved through many coordinated processes: water uptake from outside to inside the cell, subsequent pressure increase, CW creeping at the RH tip (cell expansion) and new CW synthesis(7). Thus, pressure drives tip growth and enables the RH to penetrate a mechanically resistant environment such as the soil.

Many studies focused on the growth of RH cells in liquid media under chemical or nutritive constraints(8)(9), but none were devoted to the potential effect of the mechanical resistance of the environment on the dynamics of RH growth. Thus, we decided to characterize the growth of root hair cells in mechanically resistant environments of different strength by growing RH in solid agar media of increasing concentrations (e.g. of increasing stiffnesses).

Beyond wall creep (expansion) at the RH tip, the nucleus plays an important role in RH growth in Arabidopsis thaliana (10)(11). On the one hand, it has been shown that the nucleus migrates inside the RH and is maintained at a fixed distance from the tip. On the other hand, blocking the movement of nucleus using optical tweezers was shown to reduce the growth rate of RH cells. Thus, nuclear movement and RH growth seem tightly linked. In fact, migration of the nucleus in the RH is also important for plant pathogen interaction, symbiosis, and could also be important for trichome development and male germ unit(12). Thus, investigating the sensitivity of the nucleus movement and dynamics to external stimuli is of interest for many biological processes.

Indeed, it has been reported that the nucleus responds to light(13)(14) and mechanical stimuli. For instance, by using a needle, Qu and Sun have shown that the nucleus of epidermal leaf cell is sensitive to short time and repeated mechanical stimuli(15). However, none has investigated the effect mechanical cues on the dynamics of the nucleus in growing cells such as RH. Here we characterize the effect of RH growth in media of increasing stiffnesses on the position and movement of the nucleus, a first step in the understanding of mechanotransduction in the RH.

In the following, we first investigate the effect of the mechanical resistance of the surrounding environment on the growth of RH cells. We describe the repercussions of mechanical resistance on RH growth rate, length and time of growth. To do so, we used a microfluidic-like system (MLS) that allowed us to cultivate seedling for days, and to track RH cells over up-to 3 days(16). We show that an increase in the mechanical resistance of the culture medium decreases the speed of growth and length of RH cells, but does not affect the time of growth.

Then, we describe the effect of the mechanical resistance of the culture medium on the nucleus dynamics, by looking at its speed of migration inside the RH, the fluctuations in position and speed, as well as the nucleus to tip distance. In particular, we show that the speed of the nucleus, as well as the distance between the nucleus and RH tip are reduced when the mechanical resistance of the environment is increased. These results demonstrate that the mechanical features of the environment feedback on nucleus of growing root hair cells. This is a first evidence of mechanotransduction from the cell surface to the nucleus.

## Materials and Methods

### Device preparation

The device used was prepared using a custom-made mould composed of few layers of cellulose acetate adhesive tape (3M, Magic tape) with a single channel design of 250 μm height, 1 cm width and 2 cm length. PDMS 10/1 base/curing agent mixture (Sylgard 184, Dow Corning) was poured on the mould. The PDMS was cured at 65°C for at least 4h and peeled off the mold. The PDMS chips were then bounded to glass slide using a plasma cleaner (Harrick Plasma, PDC-002-CE).

The channel was filled with warm ½ MS medium (MS Murashige and Skoog) containing 0.5% sucrose (w/w) and 0.5%, 1% or 1.25% agar (Duchefa, plant agar, w/w) PH 5.7 and a 0.5 cm thick layer of the same warm ½ MS medium was deposited on the glass slide around the PDMS chip.

### Cell Culture

*Arabidopsis thaliana’s* seeds expressing pSUN1:SUN1-GFP were used(17). After sterilization, the seeds were stratified in an Eppendorf filled with 500μL of ½ MS medium (MS Murashige and Skoog) with 0.5% sucrose (w/w) PH 5.7 for 48h at 5°C. The seeds were subsequently transferred on the device and placed on the agar medium close to the entrance of the channel. The system was then placed in a Petri dish sealed with microporous film. Afterward, the Petri dish was placed with a 45° angle with respect to the horizontal in an incubator (Sanyo, versatile environmental test chamber MLR-351H). The plantlets were kept 5 to 7 days in the incubator with a 16h light, 20.5°C and 8h dark, 17°C cycle at 65% humidity in order to wait for the root to enter the main channel.

### Microscopy / Image Acquisition

The device was placed in a microscope stage holder, few hundreds of microliter of ½ MS medium (MS Murashige and Skoog) with 0.5% sucrose (w/w) pH 5.7 were added on top of the system. Few milliliters of water were added around the system (not in contact with the agar medium) and a PVC chamber was placed above the system to limit evaporation and drying during the image acquisition. The images were recorded with a IX83 Olympus microscope equipped with a 20X0.45 NA objective. GFP fluorescence images were imaged with excitation light at 488 nm and the fluorescence emissions were recorded at 510 nm. For each plant, 42 fields of view were recorded every 10 min for 72h using a motorized scanning stage (Marzhauser SCAN IM).

### Gels Young’s modulus measurement

Cuboid (l=23mm x l=23mm x L=24mm) of ½ MS medium (MS Murashige and Skoog) containing 0.5% sucrose (w/w) and 0.5%, 1% or 1.25% agar (Duchefa, plant agar, w/w) PH 5.7 were used to measure the gels’ young modulus. The gel samples were placed between 2 glass slides. A weight m was deposited on the top glass slide and the vertical displacement Δ*L* was measured using a 5X microscope objective. The strain was kept small (~ 1%). The young modulus E of the sample was then deduced from 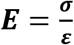 with 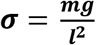 the uniform vertical stress applied on the sample, and 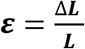 the measured strain. For each agar concentration, 5 gels were measured.

### Image analysis

To measure the physical parameters of growing root hair cells, we developed a custom-made semi-automatized pipeline using ImageJ and Matlab softwares. For each RH cell two kymographs are extracted using ImageJ, one containing the position of the tip (bright field image) and the other one containing the position of the nucleus (image of the GFP channel). Then, using Matlab, the position of the nucleus is retrieved by finding, for every time point, the maximum intensity of the GFP signal. To extract the position of the tip of the RH cell on the bright field image, we draw a polyline on the kymograph close to the tip position then the algorithm searches for the minimum intensity near this line.

### Osmotic experiments

Solid media containing ½ MS were prepared with three different agar concentrations (0.5%, 1% and 1.25%). After the autoclave step and for each concentration, two tubes of 50mL were poured with 5mL of warm medium (each of them). After cooling down to room temperature, one tube was filled with 5mL of ultrapure water and the other one with 5mL of ½ MS medium. One empty tube was filled with 5mL of ultrapure water and another one with 5mL of ½ MS medium. In order to prevent evaporation, tubes were closed with a screw cap. After 24 hours, the osmolality of the liquid phase of each tube was measured. Every experiment was repeated three times and three osmolality measurements were performed for each tube.

### Statistical analysis

Data presented are from at least 3 independent experiments. Statistical analysis was performed using either one-way ANOVA (Tukey-Kramer posthoc test) or KruskalWallis test (Dunn and Sidak posthoc test) depending on distribution properties. For normal distributions with equal variance or with at least 30 elements n (n≥30 and equal variance), one-way ANOVA tests were performed, otherwise KruskalWallis tests were performed. Distribution normality and equality of the variance were evaluated using an Anderson-Darling and a Levene’s test respectively. The statistical analysis was carried out using Matlab. p-values are reported as non-significant for p>0.05, or significant * p<0.05, ** p<0.01, *** p<0.001, **** p<0.0001.

## Results

### Length and speed of Root Hair cells, but not the time of growth, decrease when the rigidity of the medium increases

To investigate the effect of the mechanical resistance of the environment on root hair cells’ growth, *Arabidopsis thaliana* seeds were grown in agar gels of three different concentrations (0.5% w/w, 1% w/w and 1.25% w/w). At higher agar concentrations, the number of RH cells decreased drastically (1.5%w/w) or RH were even unable to grow (2%w/w, figure S1a-b). Indeed, it was recently reported in agarose gels that raising the concentration increases the force necessary to penetrate the gels (18).

In the following, we use the elastic moduli of the gels of different agar concentrations as a good proxy of their mechanical resistance to RH penetration. Thus, we measured the Young’s moduli of the gels varying agar concentration (figure S1). Young’s moduli were measured through a uniaxial compression experiment (see methods).

In order to monitor root hair cells growing for a long time period in agar gels, with a high temporal resolution, we used a microfluidic like system (MLS) that has been previously described(16). After 5 to 7 days of growth, seedlings were placed under the microscope for imaging (figure 1).

**Figure 1.**
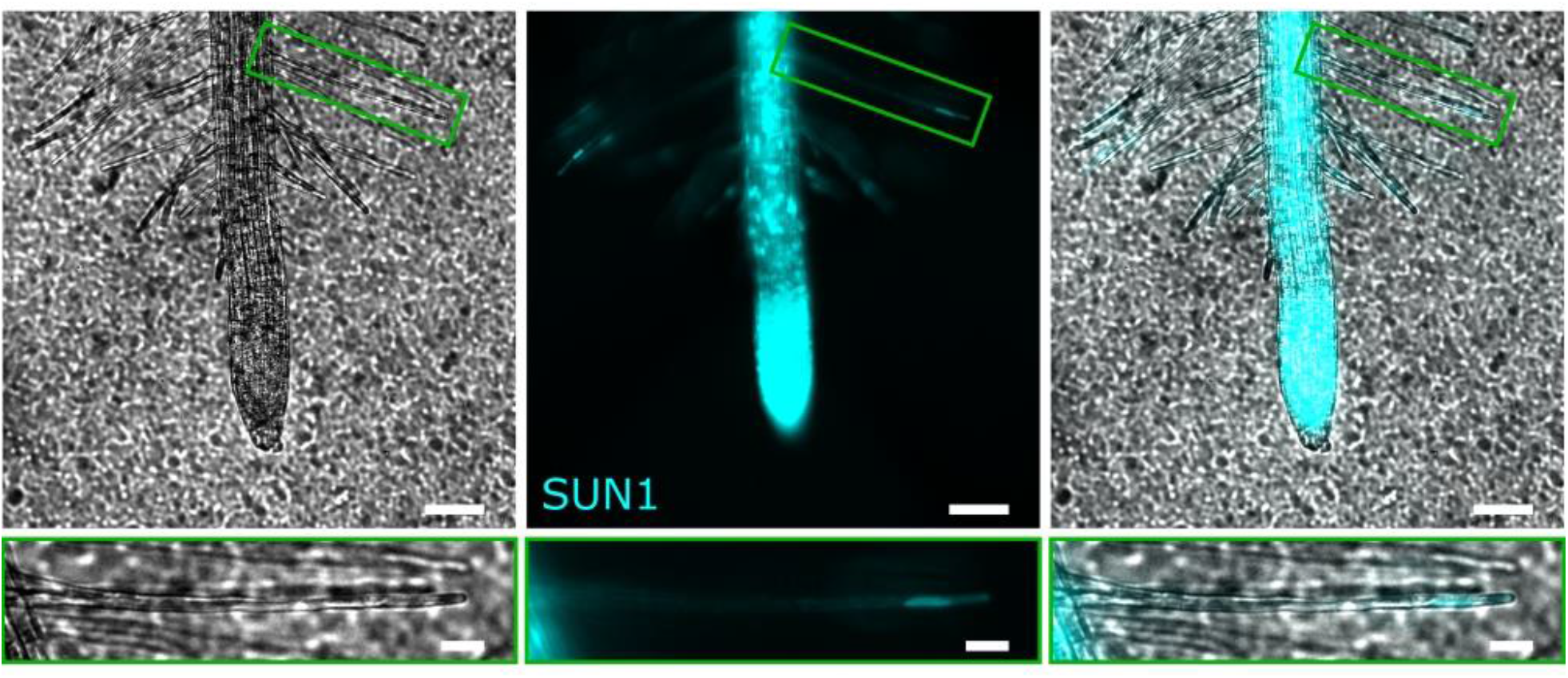
Root hair growth and nucleus imaging. Left, bright field image of Arabidopsis root and root hairs growing in ½ MS medium with 1% agar(w/w) (scale bar=100μm) and a magnification on a single root hair (scale bar =100 μm). Middle fluorescence image of pSUN1:SUN1-GFP of the same root and root hairs allowing to measure the nucleus position. Right, superposition of the brightfield and of the fluorescence images.

To reveal the effect of agar concentration on the length, speed and time of growth of RH cells, the position of the RH tip was measured over time (figure 2a) by applying a custom-made analysis pipeline on a kymograph of the Bright Field image (see methods).

**Figure 2.**
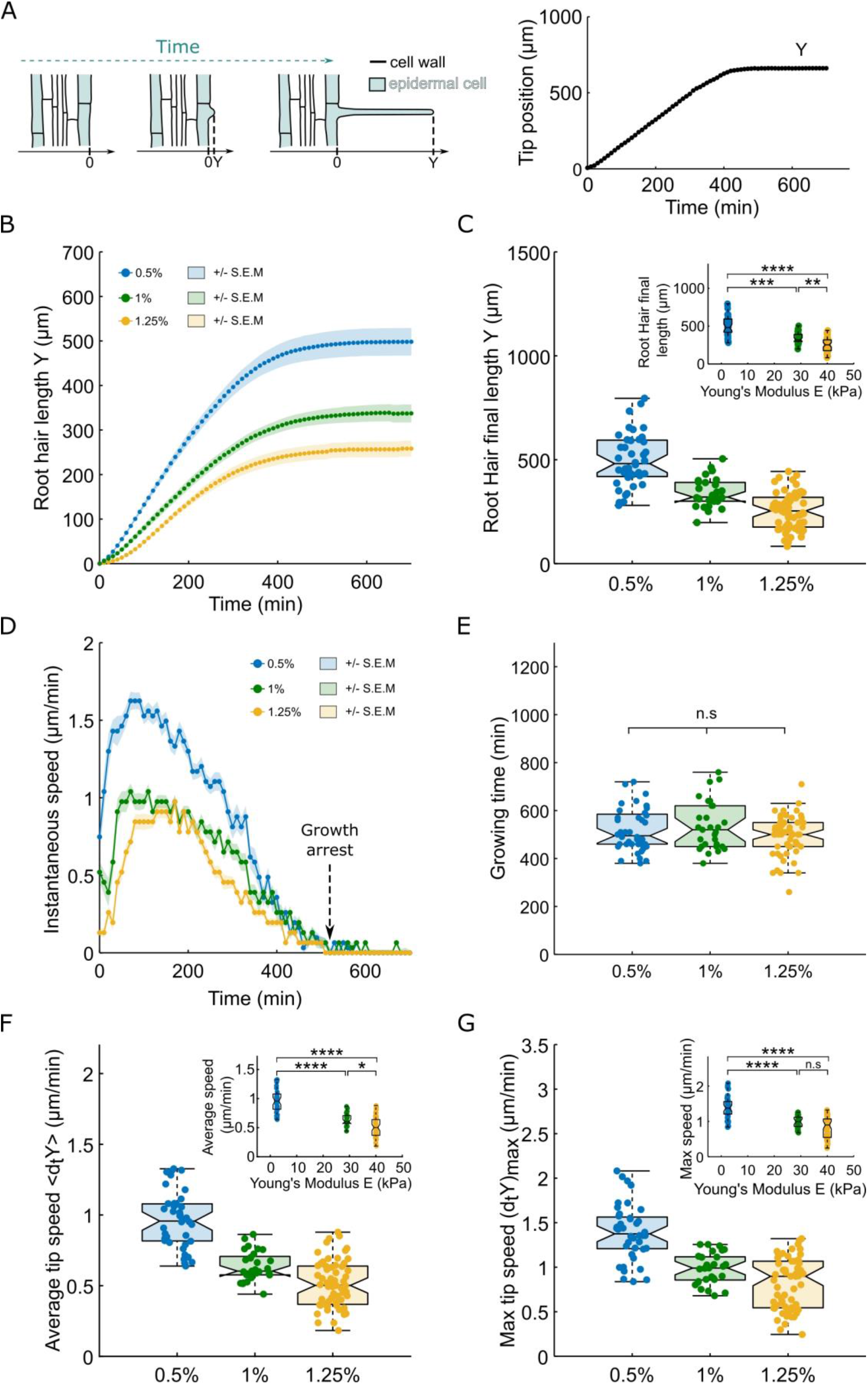
Root hair growth dynamic in agar of different concentration. (A) Left, schematic representing the growth of a root hair at 3 different times, the tip position is denoted as Y. Right, representative example of the temporal growth curve of a root hair. (B-D) Growth dynamics of root hair growing ½ MS medium with different agar concentrations, 0.5% agar (n=40), 1% agar (n=29) and 1.25% agar (n=61). (B) Average root hair growth curves +/- S.E.M. (C) Boxplots showing the distributions of the final root hair length for the 3 agar concentrations, the inset is showing the same distributions as a function of the measured agar gel’s Young’s modulus. (D) Median instantaneous growth speed curves +/- S.E.M. (E) Boxplots showing the distributions of the total root hair’s growing time for the 3 agar concentrations. (F) Boxplots showing the distributions of the average root hair tip speed during the total growth time for the 3 different agar concentrations, the insert is showing the same distributions as a function of the measured agar gel’s Young’s modulus. (G) Boxplots showing the distributions of the maximum root hair tip speed for the 3 different agar concentrations, the insert showing the same distributions as a function of the measured agar gel’s Young’s modulus.

To measure accurately the final length of living root hair cells it is important to follow them over time to avoid bias such as dead cells or growing cells (figure 2a). For each cell, we measured the length after tip growth arrest. Root hair cell length was reduced by 2-fold when agar concentration was increased from 0.5% to 1.25%. (498 +/- 20 μm, 339 +/- 12 and 253 +/- 11 μm for respectively 0.5%, 1% and 1.25%, with 40, 29 and 61 cells, figure 2 b–c). Interestingly, the median value of the RH length at 0.5% is around 500 μm, which is relatively close to the length of RH cells growing in liquid media (~580 μm)(19). This suggests that growth of RH in an agar gel of low concentration (0.5%) is similar to growth in liquid media. In contrast to the RH length, the duration of the RH growth was independent of the agar concentration, with a typical value of ~500 min (518 +/- 14 min, 535 +/- 18 and 495 +/- 10 μm for respectively 0.5%, 1% and 1.25%, with 40, 29 and 61 cells, figure 2d–e). Of note, Sun et al. reported the same ~500 min growth time in liquid media(19).

Since the RH length is reduced when agar concentration is increased, but not the time of growth, one expects the average speed of growth to decrease with the increase in agar concentration. Indeed, the analysis of the growth rates showed that the average speed of growth was reduced by 2-fold (from 0.96 +/- 0.03 μm/min to 0.51 +/- 0.02 μm/min) for agar concentrations ranging from 0.5% to 1.25% respectively (figure 2e).

A careful examination of the growth rates shows that the mechanical resistance of the environment impacts each phase of the growth, from the initiation of the RH to the regular tip growth phase. Indeed, the average speed of the RH tip during the initiation phase was drastically reduced from 1.06 +/- 0.03 μm/min to 0.48 +/- 0.02 μm/min between 0.5% and 1.25% agar gels (figure 2b–d, figure S3, figure S4). During the rapid growth phase, the maximal speed of the tip was reduced by 40% when agar concentration was varied from 0.5% (1.39 +/- 0.05 μm/min) to 1.25% (0.81 +/- 0.03 μm/min) (figure 2g). In sum, we show that the speed and length of RH cells, but not the time of growth, are reduced by the increase of the mechanical resistance of the surrounding environment. Moreover, this effect is present from the initiation phase until the end of RH growth.

Finally, in order to confirm that the effect of the agar concentration was indeed due the mechanical properties of the gels, and not due to a putative change in solutes concentrations, osmotic measurements were performed and showed no significant differences between gels of different agar concentrations (Supplementary Information and figure S2).

### Increasing the mechanical resistance of the culture medium impacts the nucleus positioning and movement in root hairs

Root hair growth and nucleus dynamics are connected(10). Since we found that RH growth depended on the mechanical resistance of the surrounding environment, we decided to determine if nuclear dynamics could also be affected. Thanks to a cell line expressing pSUN1-GFP(17), an inner nuclear membrane protein, we were able to image and track the nucleus of each cell during the whole RH growth, in gels of 0.5%, 1%, and 1.25% agar concentrations. The nucleus position over time was measured by extracting it from a kymograph of the GFP-signal (see methods).

The nucleus speed (average speed during the RH linear growth phase) is drastically reduced (1.5-fold) when agar concentration is increased from 0.5% to 1.25% (corresponding to gels Young’s moduli ranging from 2.5kPa to 40kPa) (figure 3c). In addition, the nucleus speed appeared to be nearly equal to that of the RH tip speed for the three agar concentrations tested (figure 3d).

**Figure 3.**
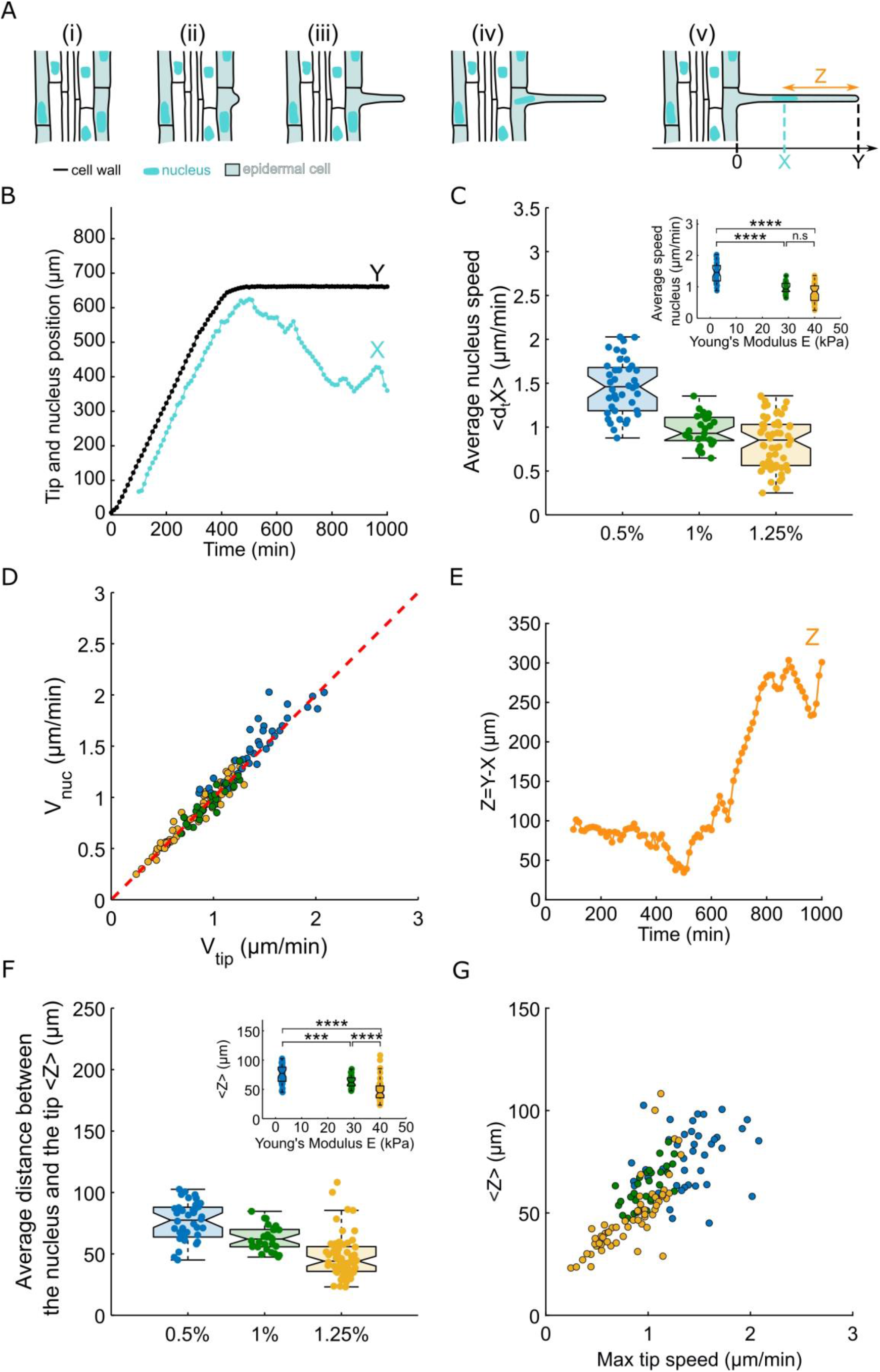
Root hair nucleus dynamic in agar of different concentration. (A) Schematics showing the growth of a root hair and the positioning of its nucleus. Y denotes the tip position and the nucleus position along the root hair is denoted as X. The nucleus position is measured as soon as the nucleus enters the root hair. (B) Representative example of the growth of a root hair and the positioning of the nucleus during growth. (C) Boxplots showing the distributions of the average nucleus speed during growth for root hair growing in ½ MS medium with different agar concentrations, 0.5% agar (n=40), 1% agar (n=29) and 1.25% agar (n=61), the inserts show the same distributions as a function of the measured agar gel’s Young modulus. (D) Graph showing the average nucleus speed as a function of the average tip speed, the red dotted line indicates where the two speeds are equal. (E) Distance between the tip and the nucleus in the example presented in (B). (F) Boxplots showing the distributions of the average distance between the nucleus and the tip for root hair growing in ½ MS medium with different agar concentrations. (G) Graph representing the average distance between the tip and the nucleus as a function of the maximum tip speed.

It has been recently described that during the RH growth the direction of migration of the nucleus fluctuates(20)(8), it has a forward mean displacement but it can go backward. Very recently, Brueggeman et al showed, by looking at the standard deviation of the speed of the nucleus, that the fluctuations of the speed increase when the microtubules are depolymerised. Looking at the standard deviation of the speed of the nucleus, we found that speed fluctuations decrease with the increase in agar concentration (figure S5 a-b). However, by normalising the standard deviation by the average speed, we found that, in proportion, the nucleus fluctuates more in a more resistant environment (figure S5c).

Interestingly, the effect of the rigidity appears at the very beginning of the RH growth, indeed, after the initiation step the nucleus entrance is delayed by the increase in rigidity. The nucleus enters in the tubular structure of the root hair after 80 minutes at low agar concentration, this time goes up to 120 minutes at high concentration (figure S6).

It has been previously described that the nucleus remains at a fixed distance from the RH tip during the fast growth phase(10).

For every RH cell, the distance Z between the tip (Y) and the nucleus (X) was computed by doing the subtraction of X from Y (figure 3a–b–c). The mean value of Z over the fast growth phase decreased from 76 +/- 2 μm at 0.5% agar concentration (soft gel of 2.5kPa) to 47 +/- 2 μm at 1.25% (stiffest gel of 40 kPa), corresponding to a 43% decrease (figure 3f). Of note, the nucleus to tip distance Z shows some subtle trends, slightly decreasing over time (figure S8).

In sum, the distance between the nucleus and the RH tip depends on the mechanical resistance of the culture medium: the stiffer is the environment the shorter is the distance between the nucleus and the tip (figure 3f). The nucleus to tip distance of 76 μm found for the 0.5% agar gels is similar to those reported previously for RH growing in liquid media(8)(10). This observation is in line with the fact that the RH growth speed in 0.5% agar gels was also similar to that reported for liquid media as mentioned in the previous section, indicating that 0.5% agar gels are soft enough to have similar impact on RH growth as liquid media.

In fact, along this line, it turns out that the rate of RH growth and the speed and positioning of the nucleus are similarly affected by the mechanical resistance of the culture medium, even quantitatively. Indeed, the distance Z between the nucleus and the tip appear to correlate well with the RH tip speed (figure 3g). This may suggest that the tip speed drives the nucleus-to-tip distance in root hair cells.

Since the RH tip growth rate and the mean speed of the nucleus are roughly the same, by calculating the ratio between the distance Z and the speed of the tip V_tip, one can have access to the lag time between the tip and the nucleus. Interestingly there is quite no effect of the mechanical resistance of the culture medium on this parameter (figure S7), the lag time is conserved.

Altogether, these results show that nuclear dynamics (speed, fluctuations, positioning) depends on the mechanical properties of the surrounding environment, probably through modulation of the RH growth rate, revealing mechanotransduction from the cell surface to the nucleus.

## Discussion

Many previous studies demonstrated the importance of external cues, such as nutrients or water availability(21)(22)(23) on the growth of root hair cells, but none looked at the effect of mechanical stimuli on root hair growth and the transduction of this signal to the nucleus. Our study represents the first step towards understanding the effect of mechanical stimuli on RH growth and their consequences on the nucleus dynamics.

The microfluidic-like system(16) allowed us to control the mechanical properties of the surrounding environment of the RH cells. In addition, we were able to follow the growth of root hair cells from the initiation step to growth arrest. Notably, we observed that increased mechanical resistance of the culture medium led to a decreased length and speed of growth of root hair cells, while the time of growth was independent of the stiffness of the medium (figure 2), indicating that reduced RH lengths are due to reduced growth rates in mechanically resisting media. Interestingly, it has been recently reported in pollen tube that the speed of growth also depends on the stiffness of agarose medium. However, depending on the species, the speed of growth can either increase or decrease when the agarose concentration increases (18).

The effect of medium mechanics on RH growth is quite different from the effect of chemical factors. For instance, Bates and lynch showed that a low phosphate availability in the medium increases the RH length and the time of growth(23). Additionally, Datta et al proposed that the transcription factor RSL4 determines the length of RH cells, the permanency of this protein is necessary to maintain the growth, when it disappears the RH cell stop to grow(22). Interestingly, the fact that in our experiments the time is constant over the three agar concentrations, make it tempting to speculate that RSL4 is not affected by mechanical cues. Thus, it would be interesting to look at presence of RSL4 in nuclei of RH cells under mechanically resistant environment.

A key component of the interaction of the root hair cell with the surrounding environment is the cell wall (CW). It is a rigid structural component that acts as a barrier to maintain cell integrity by preventing cell from bursting under osmotic constraints and protects it from damages due to mechanical stresses. The CW is composed of two layers, which are called the primary and the secondary CW. The primary cell wall (PW) is mainly composed of cellulose, hemicellulose and pectin, while the secondary cell wall (SW) is made of cellulose polysaccharides, lignin and glycoproteins. Besides the differences in composition, they have also different structures and positions/locations. The cellulose microfibrils in the PW have a random orientation, while in the SW, they are densely packed with a longitudinal alignment(24).

The PW is present on the shank and on the tip/apex of the RH cells, whereas the SW is only present on the shank. The SW rigidifies the structure which give an additional protection for the cell. During cell growth, cell wall secretion and deposition is necessary to make possible cell expansion(7). Considering that the growing time and the cell diameter are conserved but the not the length of RH cells, and if we hypothesize that the rate of cell wall deposition is the same under the different mechanical constraints, we could speculate that the cell wall thickness and/or structure could be different in highly resistant medium compare to slightly resistant medium. Thus, it will be interesting to explore the thickness, the structure and also the mechanical properties of RH cells that have grown under different medium resistances.

Thanks to a cell line expressing SUN1-GFP, we followed the dynamics of the nucleus from its entrance in the root hairs until the fully-grown step. Our experiments demonstrated the importance of the mechanical cues on the dynamics of the nucleus. As the mechanical resistance of the environment impaired drastically the RH growth, similarly, the nucleus speed was strongly reduced. Moreover, the fluctuations of the nucleus speed were modulated by the increase in mechanical resistance. Such a mechanotransduction to the nucleus was previously reported for non-growing cells, showing that a mechanical stimulus on trichome cell using a needle affects the nucleus positioning(15). They showed a quick and transient response of the trichome cell nucleus upon a short time stimulation. Interestingly, in our experiments the nucleus response starts from the initiation step by a delayed entrance in the root hair, and is observed all along the RH growth through alteration of the nucleus migration speed, of its fluctuations, and of the nucleus-to-tip distance.

Ketelaar et al have shown that the nucleus and the root hair tip are tightly linked, with a fixed nucleus-to-tip distance during RH growth. Here we show that the nucleus-to-tip-distance is dependent on mechanics, the distance being reduced for more mechanically resisting environments.

Many studies highlighted the role of actin in the link between the nucleus and the tip(11)(20)(10). It has also been reported that the nucleus dynamics is an actin and myosin-XI dependent process(25). More recently, a study suggested that during growth the backward movements of the nucleus could depend on microtubules(26).

Thus, future studies should focus on the roles of the actin and the microtubule cytoskeletons in the link between the nucleus and the RH tip, and the way they could propagate or transduce mechanical stimuli from the cell surface to the nucleus.

In conclusion, by using a custom-made microfluidic-like system, we were able to reveal the effect of the mechanical resistance of the environment on the growth of root hair cells and the subsequent alteration of nuclear dynamics and positioning. Our results represent a first step in understanding the effect of mechanical cues on the cell wall deposition, internal cell pressure regulation (e.g. on RH tip growth), as well as the transduction of the mechanical signals in root hair cells, paving the way for an integrated description and modelling of RH growth and its adaptation to mechanical properties of the environment.

## Author Contributions

DP and TA contributed equally to this work.

DP and AA developed the conceptual framework and designed the study. AA sought funding.

TA, DP and EL performed the experiments and the quantitative analyse of the results. DP, TA and AA analysed the findings and wrote the manuscript.

## Acknowledgments

We thank M-E Chabouté for fruitful discussions and for providing SUN nuclear envelope marker lines.

