## Supplementary material for "Mechanical resistance of the environment affects root hair growth and nucleus dynamics"

### 1 Supplementary Data

**Supplementary Video 1**. Root and root hair growth in agar medium. Video of the growth of a root and root hair in agar medium (1% (w/w)). Superposition of the brightfield view and, of the fluorescence signal of pSUN1:SUN1-GFP. Images were taken every 10 min.

### 2 Supplementary Information

#### 2.1 Osmotic pressure is not dependent of the agar concentration of the gels.

In order to verify that the mechanical resistance is the main parameter responsible for the observed phenotype (RH shorter, slower, etc), we measured the osmolality and calculated the osmotic pressure of the “gels” under different conditions. First, we put in contact 5mL of the gels with the same volume of ultrapure water (see methods), after 24h, we measured the osmolality of the water solution. The osmolality of the pure water was around 0 mOsm/kg of H2O, after 24h the osmolality of the water in contact with the gels was around 30 mOsm/kg of H2O for all the gels, e.g. independent of the agar concentration (figure S2a). This experiment shows that the solutes have been pumped out from the gels by osmosis, meaning that the solutes are mobile in the gels. Interestingly, this value is half of the osmolality of the liquid media used to produce the gels. Since the volumes of water and gels are equal, one should expect this value if the matric potential is negligible compared to the osmotic potential. To confirm this, another experiment was performed, instead of water, liquid media was put in contact with the gels. The osmolality of the liquid media didn’t change after 24h of contact with the gels, confirming the negligible effect of the matric potential as compared to the osmotic potential (One has to keep in mind that the concentration of MS is the same in all the gels) (figure S2b). The osmotic pressure calculated from the osmolality measurement are consistent with previous studies (figure S2b-c-)(1). Altogether, these results indicate that the reduced growth of RH and the modification of their nucleus positioning and migration for gels of increasing agar concentrations is not due to differences in water potential (matric or osmotic potential).

### 2.2 Temporal evolution of the nucleus to tip distance

If we look more precisely at the distance Z between the nucleus and the RH tip over time, we found that Z is not exactly constant during the growing phase, it decreases slowly, meaning that the nucleus comes closer to the RH tip. At the end of growth Z reaches a minimum $Z_{min}$, which is also dependent on the mechanical resistance of the surrounding environment (figure 3E, figure S8A-C). The nucleus gets closer to the tip when the resistance is high (figure S8D).

### 3 Supplementary Figures

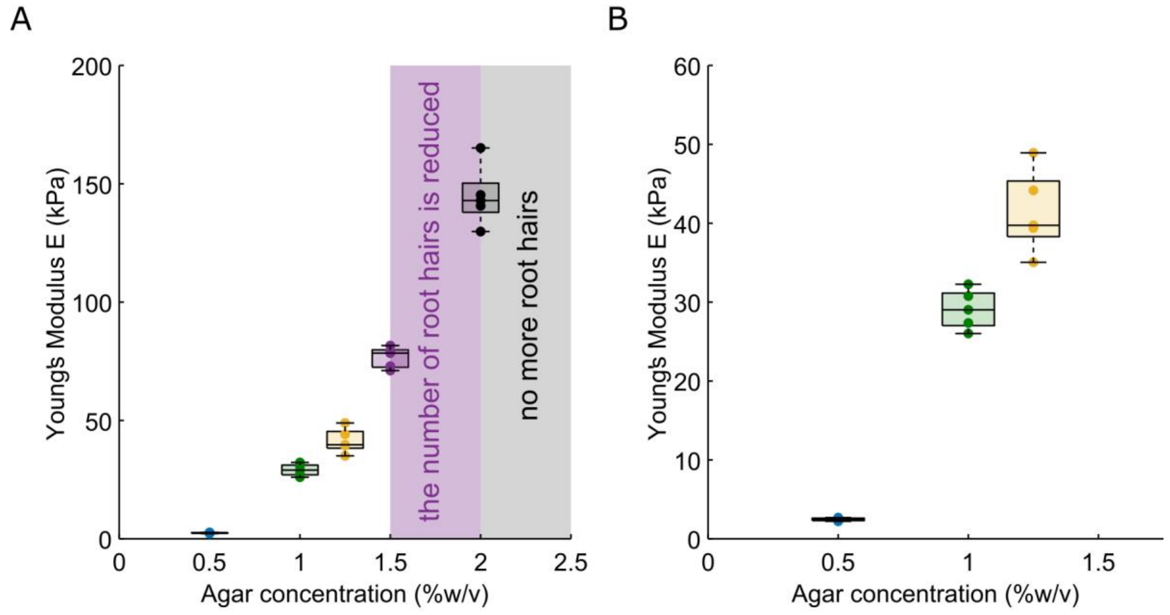


**Supplementary Figure 1.** Agar gels' Young moduli. Boxplot showing the distribution of the young modulus measured for (A) the agar concentration used in this study, (B) the agar concentration used in this study and two more agar concentration in which root hair growth is greatly impaired.

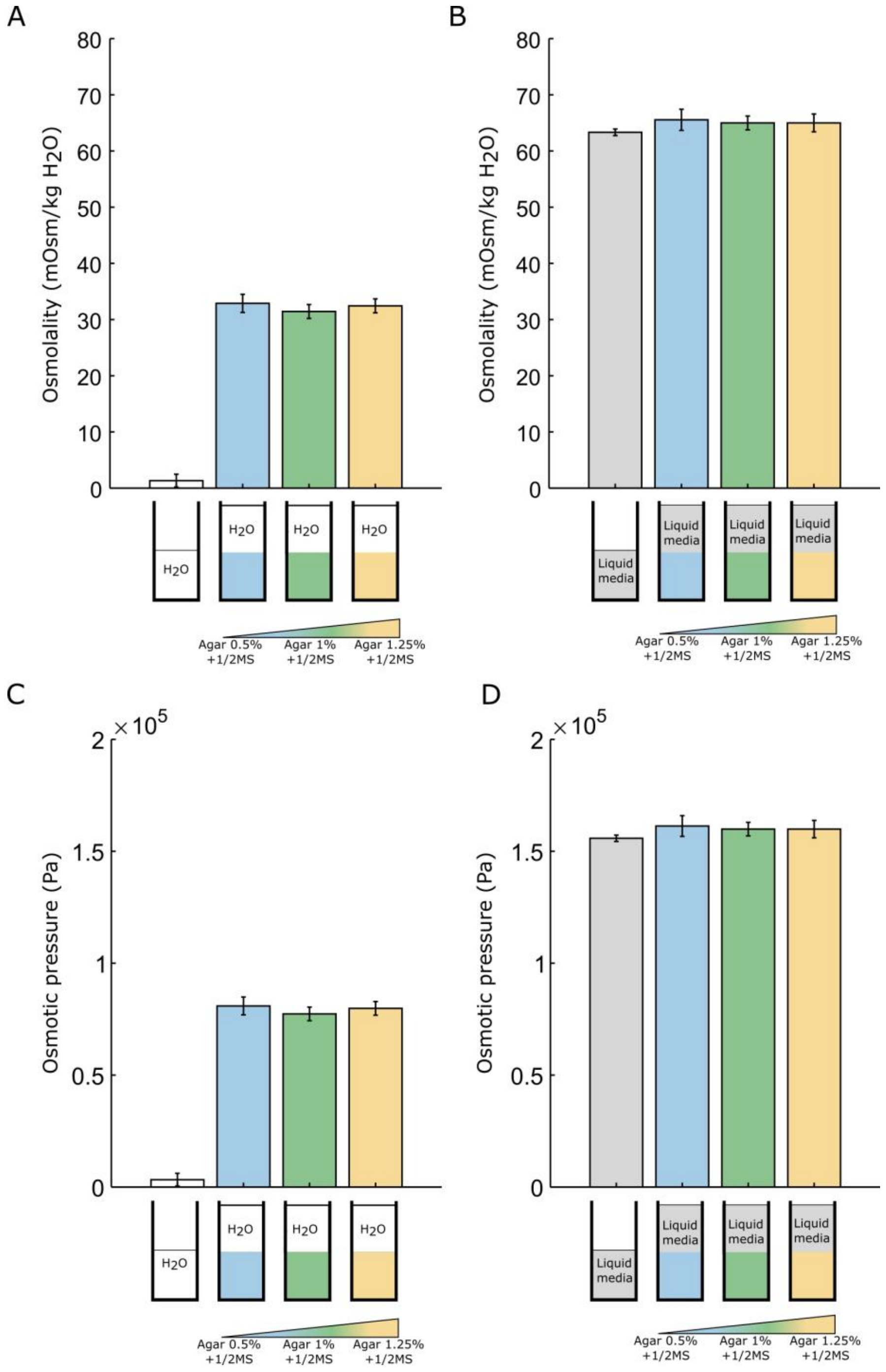


**Supplementary Figure 2.** Indirect water potential measurements of agar gels. (A) Osmolality measured for pure water and for pure water put in contact with the same volume of ½ MS agar medium with 3 different agar concentrations (0.5%, 1% and 1.25% (w/w)). (B) Osmolality measured for ½ MS medium and for ½ MS put in contact with the same volume of ½ MS agar medium with 3 different agar concentrations (0.5%, 1% and 1.25% (w/w)). (C) The Osmotic pressure corresponding to (A). (D) The Osmotic pressure corresponding to (B).

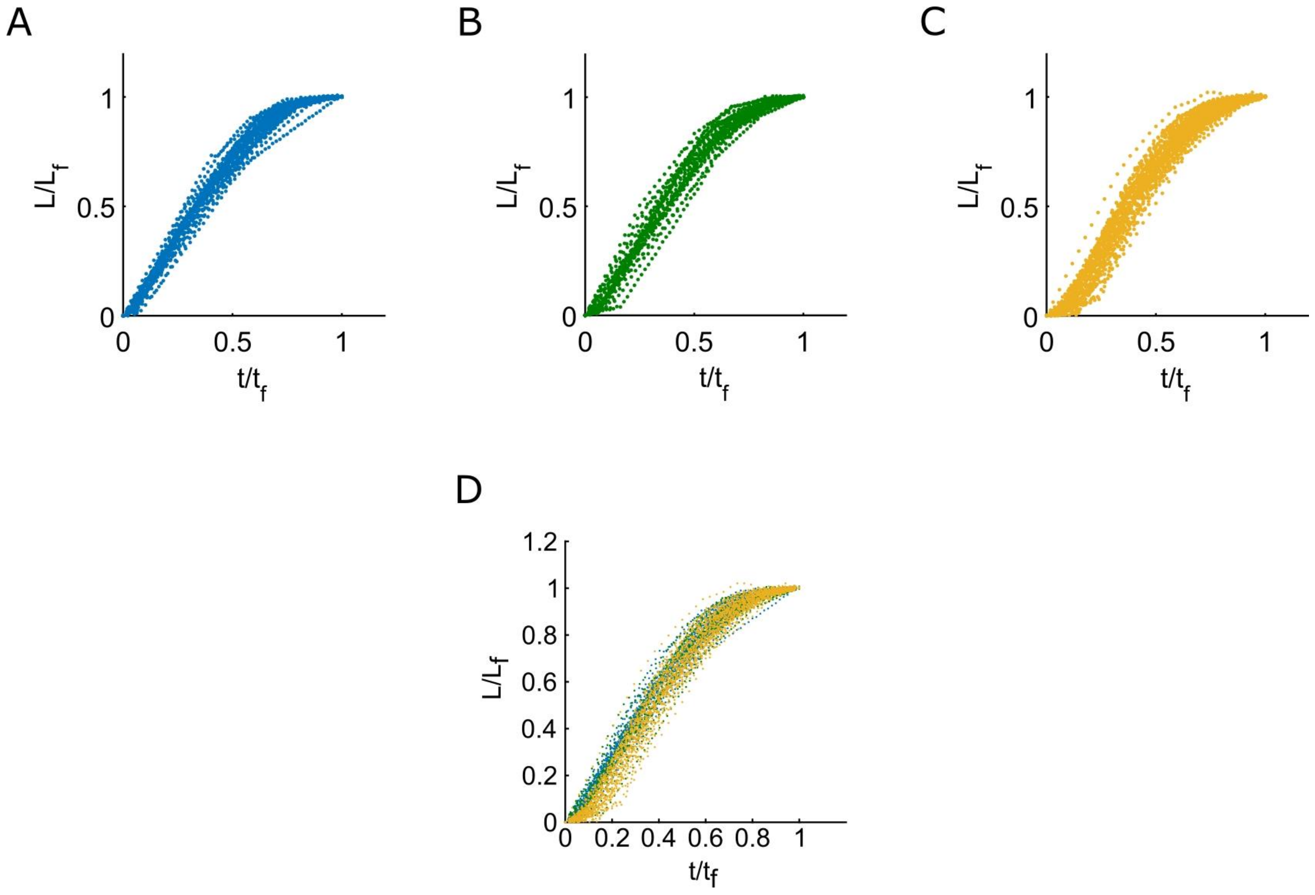


**Supplementary Figure 3.** Normalized growth curves. Superposition of normalized growth curves of root hair growing in (A) ½ MS 0.5% agar (w/w), (B) ½ MS 1% agar(w/w), (C) ½ MS 1.25% agar(w/v) and superposition of the 3 concentrations. The length is normalized by the final length and the time is normalized by the growth time.

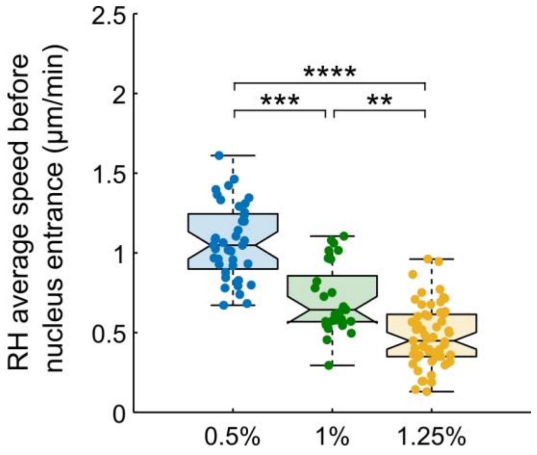


**Supplementary Figure 4.** Root hair growth speed before nucleus entrance. Boxplots showing the distributions of root hair average speed before the nucleus enters the root hair for root hairs growing in ½ MS medium with different agar concentrations, 0.5% agar (n=40), 1% agar (n=29) and 1.25% agar (n=61).

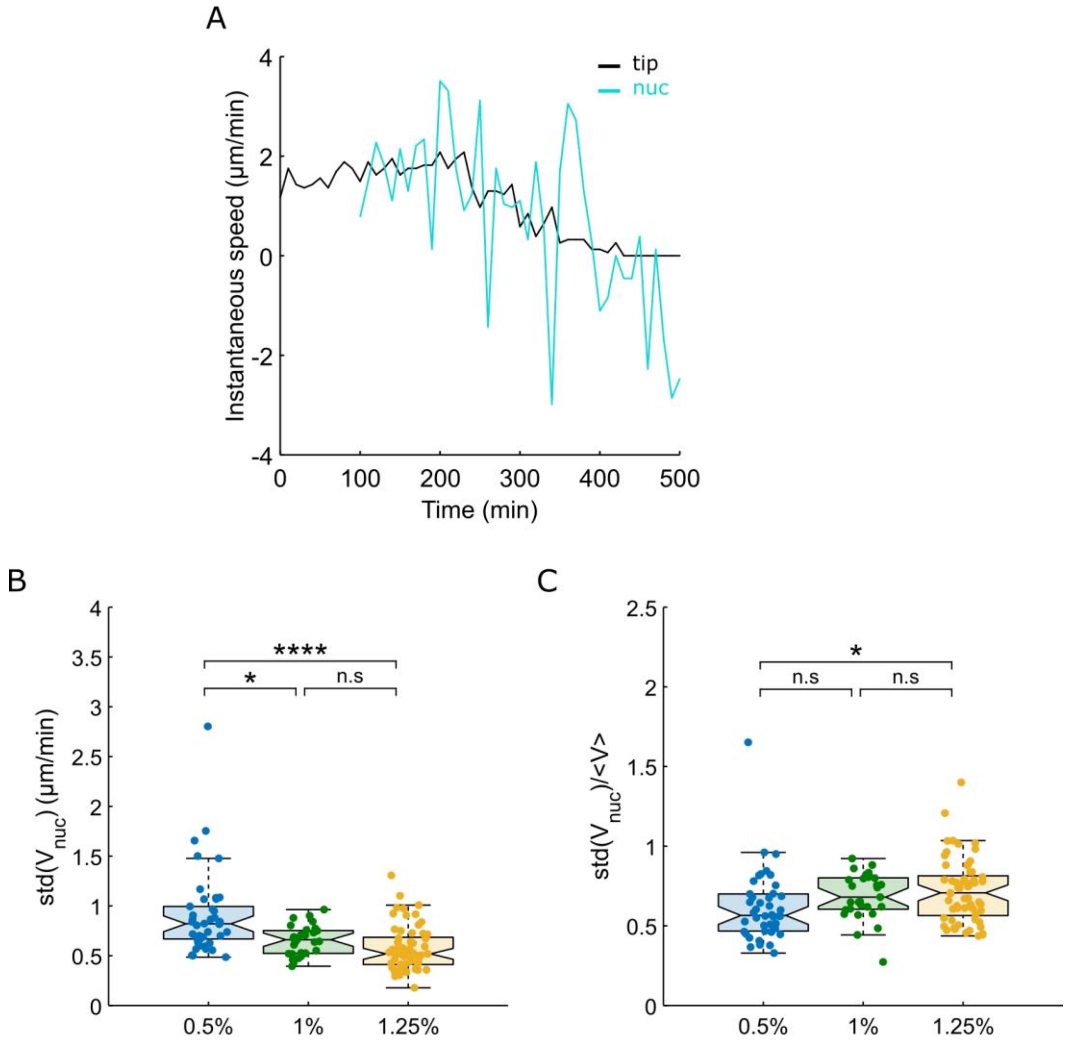


**Supplementary Figure 5.** Speed statistics. (A) Graph representing an example of the temporal evolution of the tip instantaneous speed (black curve) and the nucleus instantaneous speed (blue curve) for a root hair in a ½ MS medium with 0.5% agar concentration (w/w). (B-C) Boxplots showing the distributions of (B) the standard deviation of the nucleus instantaneous speed and of (C) The standard deviation of the nucleus speed normalized by the nucleus's average speed for root hairs growing in ½ MS medium with different agar concentrations, 0.5% agar (n=40), 1% agar (n=29) and 1.25% agar (n=61).

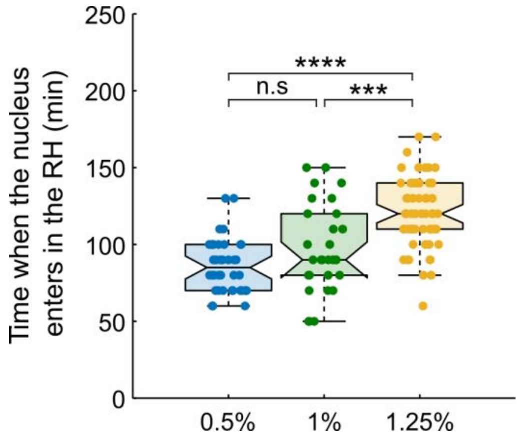


**Supplementary Figure 6.** Entry time of the nucleus in the Root hair. Boxplots showing the distributions of the time at which the nucleus enters the root hair for root hairs growing in ½ MS medium with different agar concentrations, 0.5% agar (n=40), 1% agar (n=29) and 1.25% agar (n=61).

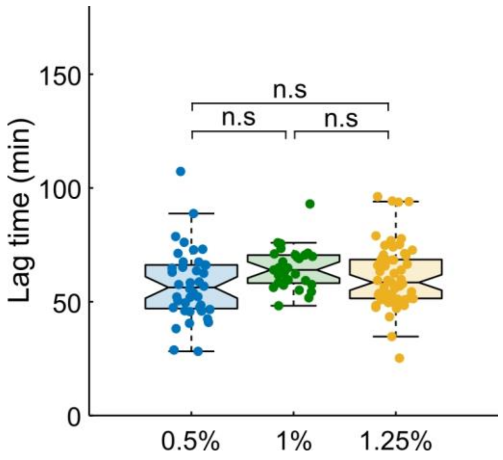


**Supplementary Figure 7.** Nucleus lag time. Boxplots showing the distributions of the nucleus lag time defined as the ratio between the average distance between the tip and the nucleus and the maximum speed of the nucleus for root hairs growing in ½ MS medium with different agar concentrations, 0.5% agar (n=40), 1% agar (n=29) and 1.25% agar (n=61).

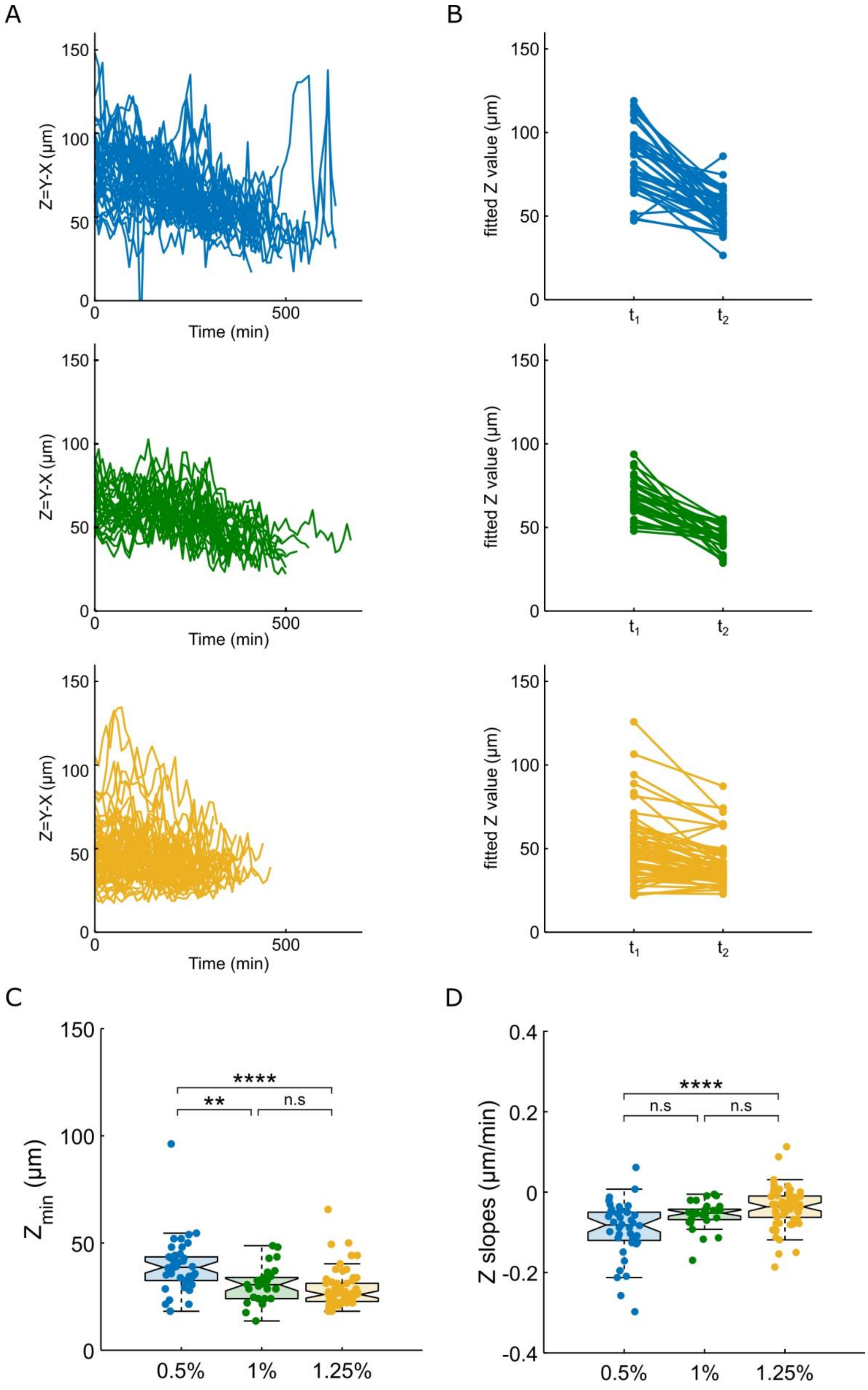

A
B
Z=Y-X (µm)
Time (min)
fitted Z value (µm)
$t_1$
$t_2$
C
D
$Z_{min}$ (µm)
Z slopes (µm/min)
0.5%
1%
1.25%
****
**
n.s

**Supplementary Figure 8.** Nucleus dynamics. (A) Superposition of the curves of the distance between the tip and the nucleus from the the beginning of the linear phase of growth to the time corresponding to the minimum distance between the tip and the nucleus for root hairs growing in ½ MS with 0.5% agar (top) or with 1% agar (middle) or with 1.25% agar (bottom). The minimum is taken as the minimum for +/- 100 minutes around the growth arrest. (B) Superposition of the linear fit of each curve of (A) with t1 the beginning of the linear phase of growth and t2 the time corresponding to the minimum distance between the tip and the nucleus. (C) Boxplots showing the distributions of the minimum distance between the tip and the nucleus for root hairs growing in ½ MS medium with different agar concentrations, 0.5% agar (n=40), 1% agar (n=29) and 1.25% agar (n=61). The minimum is taken as the minimum for +/- 100 minutes around the growth arrest. (D) Boxplots showing the distributions of the slope of the linear fit of each curve in (A).
